# The two groups of zebrafish type I interferons target different tissues, paralleling the mammalian type I: type III IFN functional division

**DOI:** 10.1101/2025.11.03.686239

**Authors:** Hannah Wiggett, Dean Porter, Valerie Briolat, Teodosia Balaboi, Maroun Abi Younes, Ingrid Colin, Georges Lutfalla, Jean-Pierre Levraud

## Abstract

Interferons (IFNs) are ancient cytokines that arose in jawed vertebrates ∼400-500 million years ago. IFN systems are present with conserved antiviral functions across vertebrate lineages, including zebrafish (*Danio rerio*). In mammals, antiviral IFNs are divided between type I interferons (IFN-I), which drive systemic responses, and type III interferons (IFN-III), which protect barrier mucosal epithelia, owing to the specific distribution of their respective receptors. Although zebrafish lack IFN-III, they have IFN-Is which subdivide into 2 groups with distinct receptors, providing a unique opportunity to study how antiviral immunity has evolved in the absence of IFN-III. Whilst previous work has suggested complementary, non-redundant roles for IFNs from these groups, the tissue specificity has not yet been resolved.

As larvae, zebrafish only express one group 1 (IFNφ1) and one group 2 IFN (IFNφ3). Using viral infection assays and reporter transgenics, we found that IFNs from group 1 (IFNφ1) and group 2 (IFNφ3) are produced by distinct subsets of cells, with no detectable co-expression. To assess tissue and cell-type-specific responses to these two IFNs, we used ISG reporter fish imaging and whole-larva single cell RNA sequencing after injection of recombinant IFNφ1 and IFNφ3. Despite a similar core ISG response, distinct downstream ISG programs across multiple tissues and organ systems were found. In particular, barrier epithelial cells, such as enterocytes, responded more strongly to IFNφ1, while myeloid cells responded more strongly to IFNφ3. Our results indicate that zebrafish IFN-I families have functionally diversified their antiviral immune responses by tissue context, driven by cellular partitioning of both IFN-I production and response. These results mirror the division of labour between mammalian IFN-I and IFN-III, emphasising the evolutionary importance of tissue division of immune responses, as well as deepening our understanding of the zebrafish as a model for host-pathogen interactions.

## Introduction

Interferons (IFNs) constitute the major cytokine family induced by viral infection. In mammals, antiviral defence is primarily coordinated by type I (IFN-I) and type III IFNs (IFN-IIIs otherwise known as IFN*λ*), which both activate canonical JAK-STAT pathways and drive the transcription of overlapping sets of interferon-stimulated genes (ISGs) (Lazear et al. 2019; Zhou et al. 2007). The critical functional distinction between IFN-I and IFN-IIIs lies in their cell type specificity which is determined by differential receptor usage. IFN-Is signal broadly, with almost all cell types expressing their cognate IFNAR1/2 receptor. In contrast IFN-IIIs are cell type specific with their receptor, consisting of IFNLR in complex with IL10RB, largely limited to mucosal barrier epithelial tissues, such as the respiratory and gastrointestinal tracts (Pestka et al. 2004; Sommereyns et al. 2008). This receptor distribution underlies their complementary roles with IFN-IIIs providing a front-line defence at epithelial barriers, while IFN-Is result in broader systemic antiviral immunity.

Despite extensive progress, several challenges limit our understanding of IFN biology. *In vitro* systems have shown that IFN responses are highly context-dependent, influenced by factors such as receptor density, cell polarity, and epigenetic state (Bhushal et al. 2017; Pervolaraki et al. 2018; Mostafavi et al. 2016). Moreover, clinical applications of IFNs are limited by off-target effects and tissue-specific toxicity, reflecting the difficulty in predicting IFN responses outside of the context of the whole organism. This has led to a growing emphasis on studying IFNs *in vivo*, where intercellular communication, developmental context, and tissue organisation can be resolved.

Zebrafish (*Danio rerio*) are small vertebrate models, which have now been established as valuable models for host-pathogen interactions (H. Meijer and P. Spaink 2011; Levraud et al. 2014; Sullivan et al. 2021). Their small size, optical transparency and availability of transgenics offer the ability to track organism wide responses with high resolution. Importantly zebrafish also possess dedicated antiviral IFNs, which originated ∼400–500 million years ago in early jawed vertebrates (Redmond et al. 2019).

Phylogenetic studies have now identified what appear to be IFN-IIIs in cartilaginous fishes (Redmond et al. 2019), however in zebrafish there are no IFN-IIIs, despite early claims (Levraud et al. 2007) which were refuted by structural analysis (Hamming et al. 2011). Instead, zebrafish possess 4 antiviral IFN-I genes (IFN-φ1 to IFN-φ4) which have been shown to confer protection against a range of human and fish viruses (Laghi et al. 2024; Ge et al. 2015; Palha et al. 2013; Boucontet et al. 2018; Aggad et al. 2009; López-Muñoz, F. Roca, et al. 2009). Of note, recently a new type of IFNs, type IV IFNs, have been identified with IFN-ν in zebrafish (Chen et al. 2022) and in other lineages (Deng et al. 2025) but its functional importance remains to be established.

Broadly, fish IFN-Is are classified into two groups based on the number of their disulphide bridges (Zou et al. 2007). In zebrafish, these two groups have separate receptors that have one specific and one shared chain, with group 1 IFN-Is (IFNφ1 and IFNφ4) binding to a CRFB1/CRFB5 dimer; while group 2 IFNs (IFNφ2 and IFNφ3) bind to a CRFB2/CRFB5 dimer (of note, due to the incorrect initial assumption that zebrafish IFNs were orthologous to IFN-IIIs, the current official name of crfb5 is il10rb, we will use instead crfb5 here). Structural analyses confirmed that both IFN groups are homologous to mammalian IFN-Is (Hamming et al. 2011), leading to the hypothesis that the two groups arose during the teleost specific whole genome duplication (WGD). Interestingly, such duplication events are important drivers for evolution, helping to give rise to genes with new functions (Ohno 1999). Genomic analysis indicates a more complex situation, with probable diversification via tandem duplication predating the teleost WGD (Boudinot et al. 2016; Liu et al. 2019). Nevertheless, distinguishing between redundancy, sub-functionalisation or the development of new functions amongst paralogous genes can be useful to inform us of evolutionary trajectory (Kuzmin et al. 2021).

In the adult zebrafish, IFNφ1 and IFNφ3 play complementary roles, with distinct kinetics (López-Muñoz, F. J. Roca, et al. 2009) while work conducted *in vivo* (Aggad et al. 2009) or using cell lines has shown that IFNφ4 has very little antiviral capacity compared to IFNφ1 (Chen et al. 2024). These observations suggest functional diversification exists within the zebrafish IFN system, yet there has been very little work done to understand the cell-type and tissue-specific distinctions between IFN-I groups. Although larval zebrafish already display a strong IFN-I response, they only express IFNφ1 (group 1) and IFNφ3 (group 2) to significant levels (Aggad et al. 2009; Briolat et al. 2014), providing a simplified setting to address this question. Here, we address this gap by characterising the spatial and transcriptional differences between IFNφ1 and IFNφ3 in larval zebrafish, providing insight into their functional divergence and highlighting parallels between mammalian IFN-I and IFN-III systems.

## Results

### IFNφ1 and 3 induce comparable but distinct ISG expression profiles

To first ensure a sufficient and comparable ISG response to IFNφ1 and IFNφ3, we used transgenic larvae expressing GFP and mCherry under the control of isg15 and mxa promoters respectively (Figure 1A). These two lines also harbour secondary fluorescent reporters in the lens and the heart, respectively, but for simplicity will be designated as Tg(mxa:mCherry) and Tg(isg15:eGFP). To induce an organism wide IFN response, we injected 4 day post fertilisation (dpf) larvae with recombinant IFNφ1 and IFNφ3 into both the coelomic cavity and the brain. Low magnification imaging of the whole larvae demonstrated that injection of either IFNφ1 or IFNφ3 resulted in robust induction of fluorescence for both reporter lines at 24hpi (Figure 1B). Flow cytometry of the whole dissociated larvae confirmed this, showing slight differences between isg15 and mxa reporters. Both IFNs induced the expression of the mxa reporter but with a larger proportion of cells responding to IFNφ3 than IFNφ1 (Figure 1C), albeit with similar median fluorescence intensities (Figure 1D). In contrast, neither the proportion of cells expressing the isg15 reporter (Figure 1E) or the median fluorescence intensity of these cells was significantly different between IFNs (Figure 1F).

**Figure 1.**
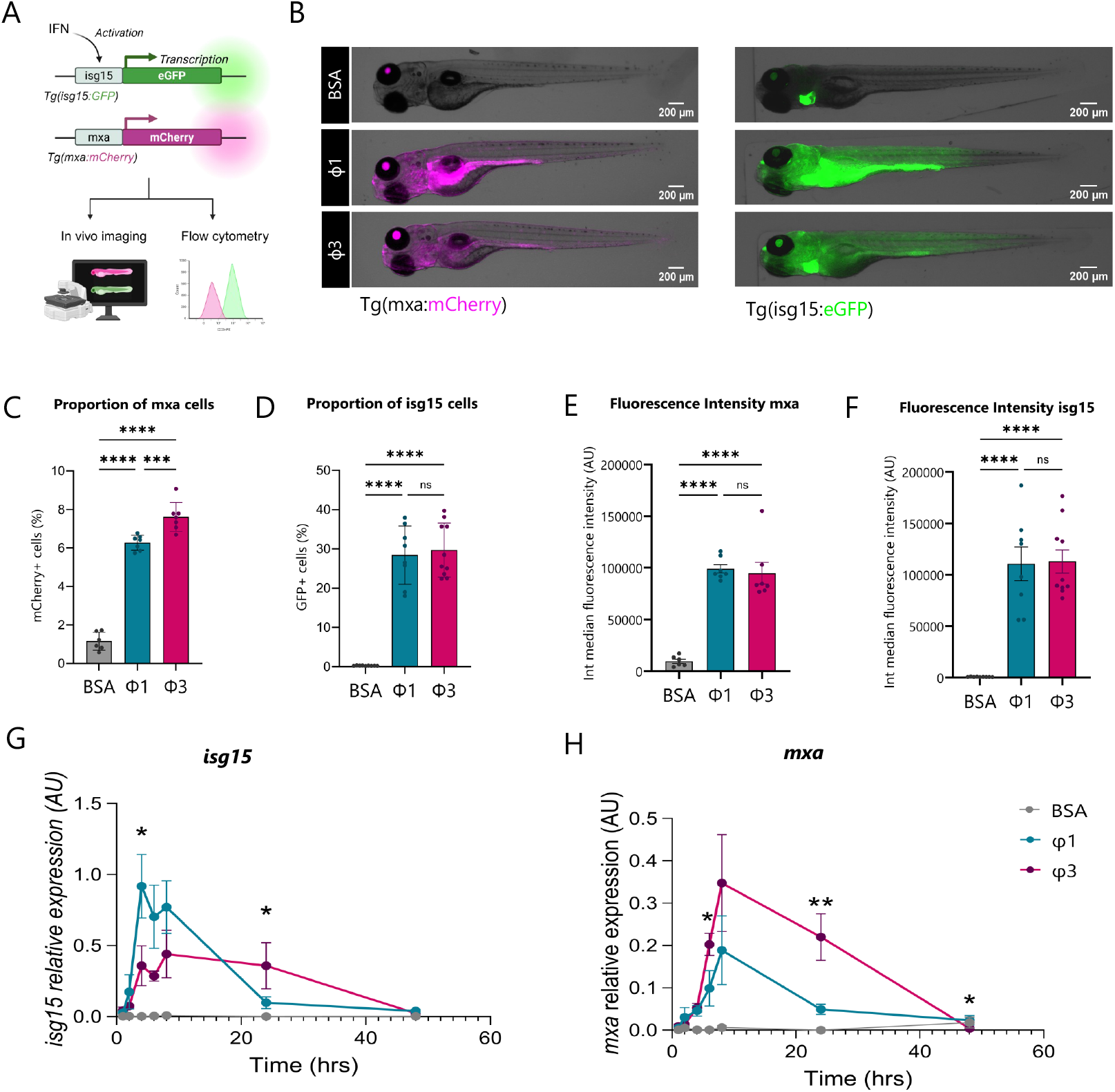
Global ISG induction by IFNφ1 and IFNφ3. **A**. Transgenes used for mxa and isg15 expression by microscopy and flow cytometry **B**. Widefield images of whole Tg(isg15:GFP) and Tg(mxa:mCherry) larvae 24hpi with rIFNφ1 and rIFNφ3. Images are single plane. GFP Fluorescence in the heart and mCherry in the eyes are transgene markers. **C**. Proportion and **D**. integrated median fluorescence intensity of mCh+ cells in Tg(mxa:mCherry) and **E. F**. Tg(isg15:GFP) larvae 24hpi of rIFNφ1 and rIFNφ3 or control (BSA) in 4dpf zebrafish larvae (n = 7-11 per condition, pools of 3 larvae per replicate). Data were tested for significance using one way ANOVA with multiple compari-sons. Error bars represent the SEM and asterisks indicate statistical significance **p<0.01, ****p<0.0001. **G**. rtqPCR of isg15 and **H**. mxa, transcript expression relative to EF1a following coelomic and intracerebral injection of rIFNφ1 and rIFNφ3 or control (BSA) in 4dpf zebrafish larvae (n=5-8 per time point, pools of 3 larvae per replicate). Data were tested for lognormal distribution using GraphPad prism and comparisons between φ1 and φ3 were made using lognormal Welch’s t test to account for unequal variances. Error bars represent the SEM and asterisks indicate statistical significance *p < 0.05, **p<0.01. Only statistical tests between φ1 and φ3 are shown, for statistical tests across all time points see supplementary table 1.

To then investigate the kinetics of the ISG responses, we performed qPCRs to examine targeted gene expression over the course of 48 hours. Analysis showed that transcription of *isg15* (Figure 1G) was increased at all time points following injection with both IFNφ1 and IFNφ3 compared with BSA-injected controls and at all time points between 4 and 24 hours for *mxa* (Figure 1H) (Supplementary Table 1). Some differences in the kinetics were observed, *isg15* expression peaked at 4 hours post injection (hpi) in response to IFNφ1 injection, whereas IFNφ3 peaked at 8hpi. The response after IFNφ1 injection was also faster but transient, whereas expression was maintained at 24hpi after IFNφ3 injection. *isg15* and *mxa* showed markedly different transcriptional patterns, with IFNφ3 inducing stronger transcription in *mxa* and IFNφ1 in isg15. The peak of expression in response to *mxa* was at 8hpi for both IFNφ1 and IFNφ3, but was stronger in response to IFNφ3, which showed significantly increased expression of *mxa* compared with IFNφ1 at both 6 and 24hpi. Despite heterogeneity between ISGs, the administered doses produced strong, comparable ISG induction, ensuring validity of comparisons of IFNφ1 and IFNφ3.

**Table 1.**
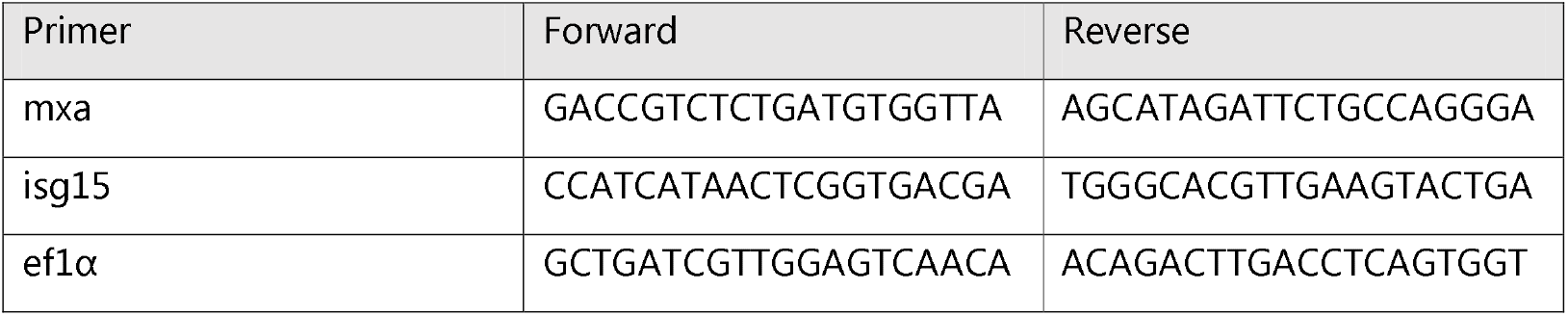
Primer sequences used for qPCR.

### IFNφ1 and IFNφ3 activate shared antiviral programs with distinct signatures

To further examine the differences in ISG responses at the cellular level we performed single cell transcriptomics on whole larvae injected with recombinant IFNφ1, IFNφ 3 or BSA as a control (Figure 2A). To ensure capture of the direct transcriptional response, samples were analysed at 4hpi, a timepoint shown to represent one of the earliest windows of detectable transcriptional responses following immune stimulation (Mostafavi et al. 2016; Lee et al. 2025; Cui et al. 2024).

**Figure 2.**
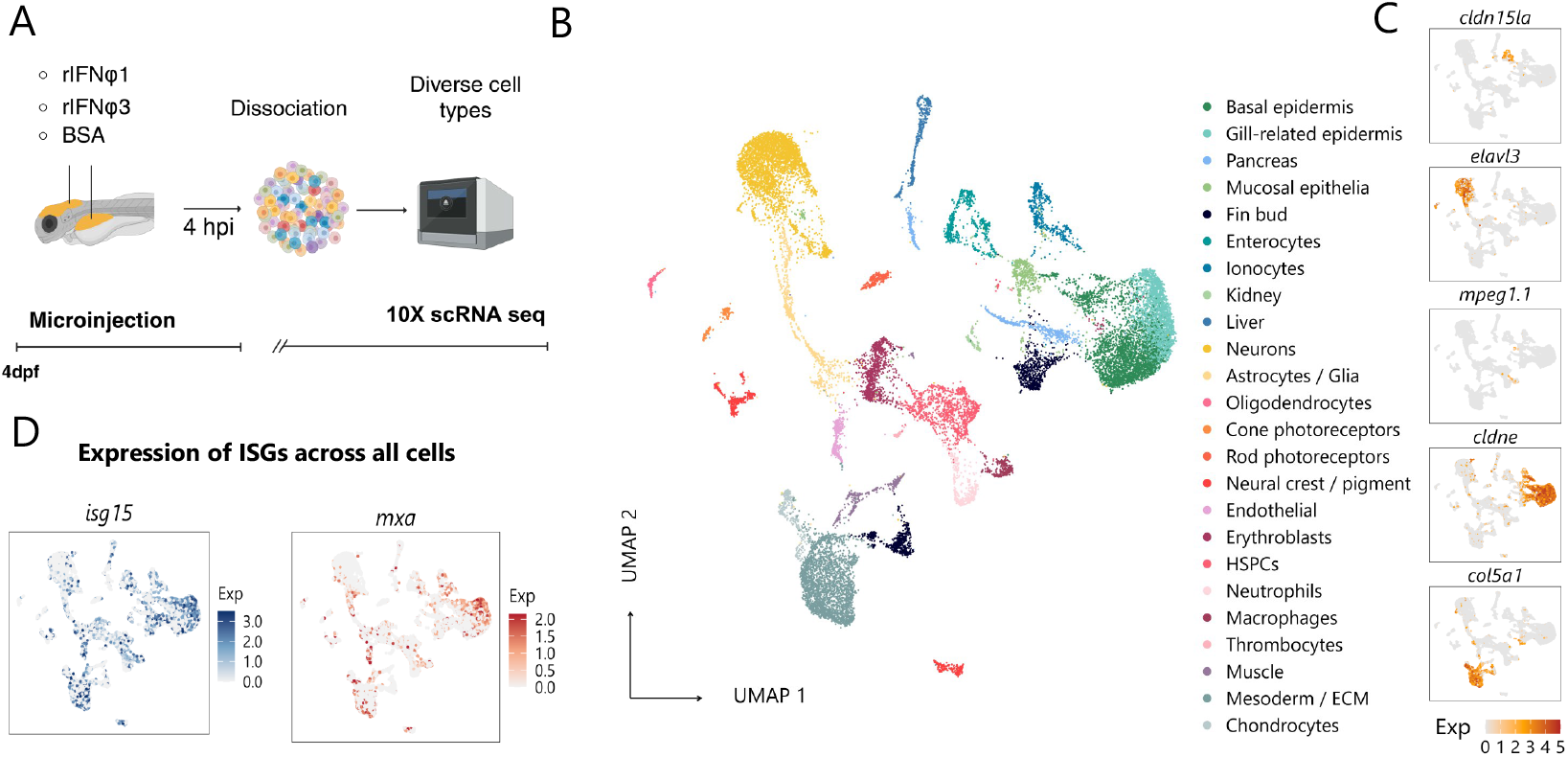
Whole larvae transcriptomics of ISG responses. **A**. Schematic representation of experimental design: 4h post intracerebral and coelomic cavity injection of recombinant IFNs or BSA (control), whole Tg(elavl3:RFP;gfap:GFP) larvae were dis-sociated (12-15 pooled larvae per treatment condition) and 10X single cell sequencing librar-ies were prepared **B**. UMAP projection of clusters obtained after QCs and removal of erythro-cyte clusters and doublets. Dimensional reduction was performed using PCA on the top 2,000 variable genes (25 PCs). Clustering was performed using the Louvain algorithm at resolution 0.6. **C**. Feature plots displaying log-normalised expression of canonical marker genes used for cell type annotation: *cldn15la* (enterocytes), *elavl3* (neurons), *mpeg1*.*1* (macrophages), *cldne* (epidermis), and *col5a1* (mesoderm). Colour intensity reflects log-normalised expression per cell (see legend). **D**. Bar plots showing the fraction of each annotated cell type across treatment conditions (BSA, IFNφ1, IFNφ3). Proportions are expressed as the fraction of cells within each group **E**. Feature plots illustrating log-normalised expression of canonical ISGs (*isg15, mxa, ifit14*) across the UMAP. Colour intensity reflects log-normalised expression per cell (see legend).

After quality controls and filtering to exclude nucleated erythrocytes which respond very weakly to IFNs, we obtained transcriptomes from 22,341 cells, from which we identified 24 clusters from diverse types (Figure 2B). Clusters were annotated manually according to known marker expression and comparison with the Farnsworth atlas for the developing zebrafish (Farnsworth et al. 2020) (Figure 2C, Supplementary Figure 1, Supplementary Table 2). All clusters contained cells from each treatment group (Supplementary Figure 2) and across the dataset, spatial mapping of *isg15* and *mxa* transcript expression on the umap localises their expression with cellular resolution, with IFN-induced signal detectable in every annotated cell type (Figure 2D).

Before dissecting the cell type specificities of the response, we used pseudobulk differential gene expression (DEG) analysis of treatment groups compared with controls to identify transcriptional changes present amongst all cells in response to IFNs. This demonstrated the robust upregulation of ISGs in response to both IFNs (Figure 3 A, B). The majority of the differentially expressed genes were upregulated rather than downregulated. Most upregulated DEGs were shared between the two IFNs and consisted primarily of canonical ISGs including *mxa* and *isg15* as well as several members of the *ifit* and *trim* families, among others (Figure 3C), consistent with our previous bulk analysis of response to IFNφ1 (Levraud et al. 2019). It also included previously reported unannotated zebrafish genes which have been reported to be upregulated during viral infection such as CU984600.2 (Hu et al. 2024).

**Figure 3.**
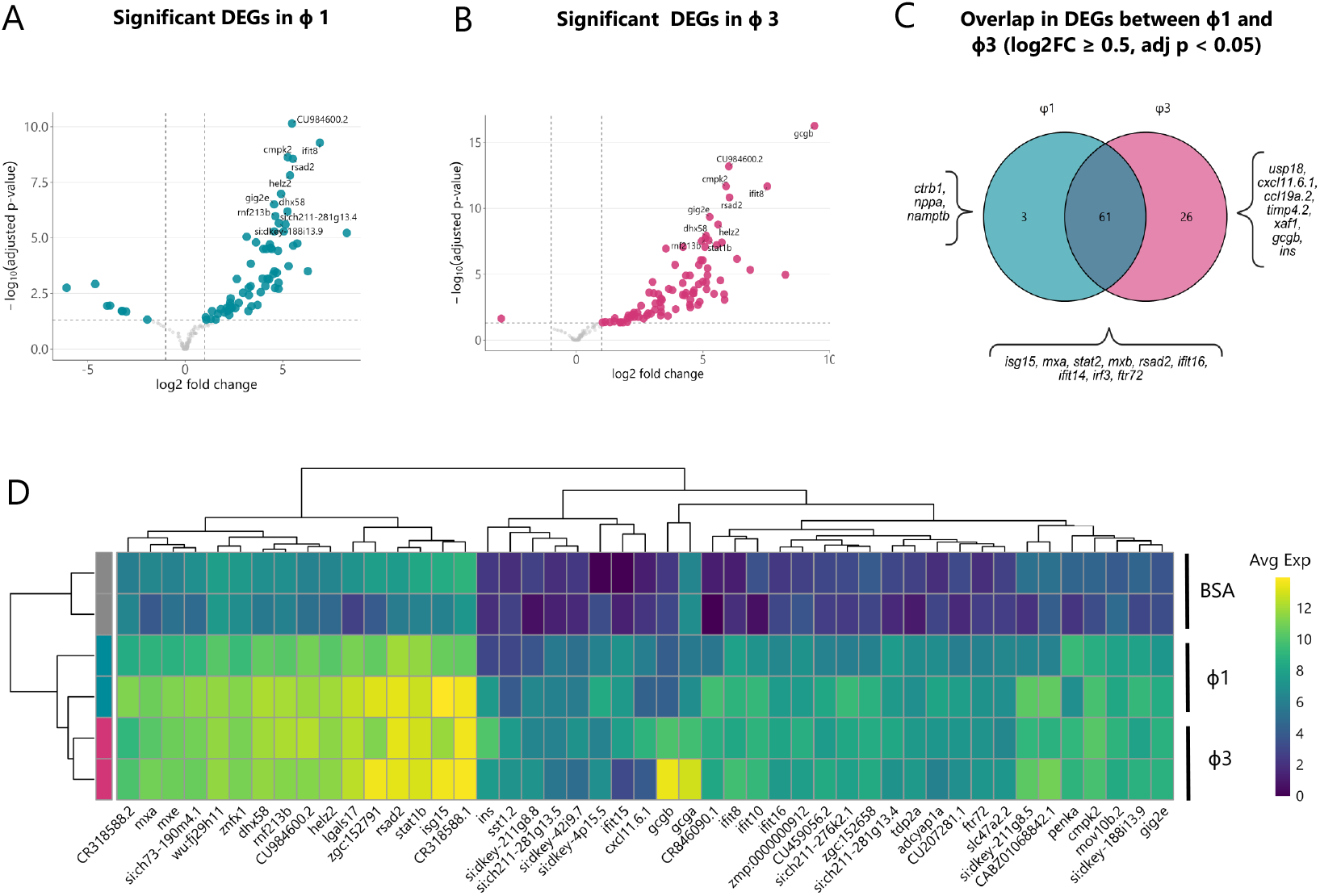
Pseudobulk differential expression analysis of organism wide transcriptional responses. Volcano plots for **A**. IFNφ1 vs BSA and **B**. IFNφ3 vs BSA. The x-axis shows log2fold change and Y axis the log-10 adjusted p values. Points are coloured by significance upregulated (p adj. < 0.05 and log2FC > 0.3) and downregulated (p adj. < 0.05 and log2FC < −0.3). Horizontal and vertical reference lines mark the FDR and fold change cut-offs. Top genes are labelled. Differential expression was computed using DESeq 2, FDR values are Benjamini-Hochberg adjusted. n=2 independent replicates (12—15 pooled lar-vae each) per treatment group. **C**. Venn diagram showing overlap in significantly upregulated (p adj. < 0.05 and log2FC < 0.3) DEGs between conditions **D**. Heatmap of log 2 normalised average expression per sample for the union of the top 40 upregulated DEGs in IFNφ1 and IFNφ3. Rows represent the samples, n = 2 per treatment group as above. Both rows and columns are hierarchically clustered using Euclidean distance and complete linkage.

Some differences were seen, mostly within genes expressed at lower fold changes. Globally, IFNφ3 upregulated a larger spectrum of genes with 26 upregulated DEGs unique to this condition. Of note, these included immune genes such as C-X-C motif chemokine 11 (*cxcl11*.*6*.*1*) and chemokine ligand 9 (*ccl19a*.*2*) and genes involved in glucose metabolism, such as glucagon b (*gcgb*) and insulin (*ins*). IFNφ1 showed a much narrower unique response, with only three upregulated DEGs, Chymotrpsinogen B1 (*ctrb1*) a digestive protease precursor, nicotinamide phosphoribosyltransferase (*namptb*) an NAD+ salvage pathway enzyme and the secreted hormone natriuretic peptide A (*nppa*). Despite these differences, plotting of the most significantly expressed genes in either condition revealed globally similar transcriptional patterns to both IFNs (Figure 3D).

### IFN-φ3 induces broader transcriptional and antiviral programs across more tissues than IFN-φ1

To dissect cell-type specific responses to each IFN, we performed single-cell level DEG analysis within each annotated cell type. To ensure that the number of DEGs between treatment groups was unbiased from cell number, we down-sampled by cell type prior to DEG analysis. The transcriptomic response to IFN-φ3 tended to be broader, resulting in the upregulation of more DEGs than IFN-φ1 in most cell types, with the exception of the gill-related epidermis, Mesoderm/ECM, fin bud and neurons (Figure 4A).

**Figure 4.**
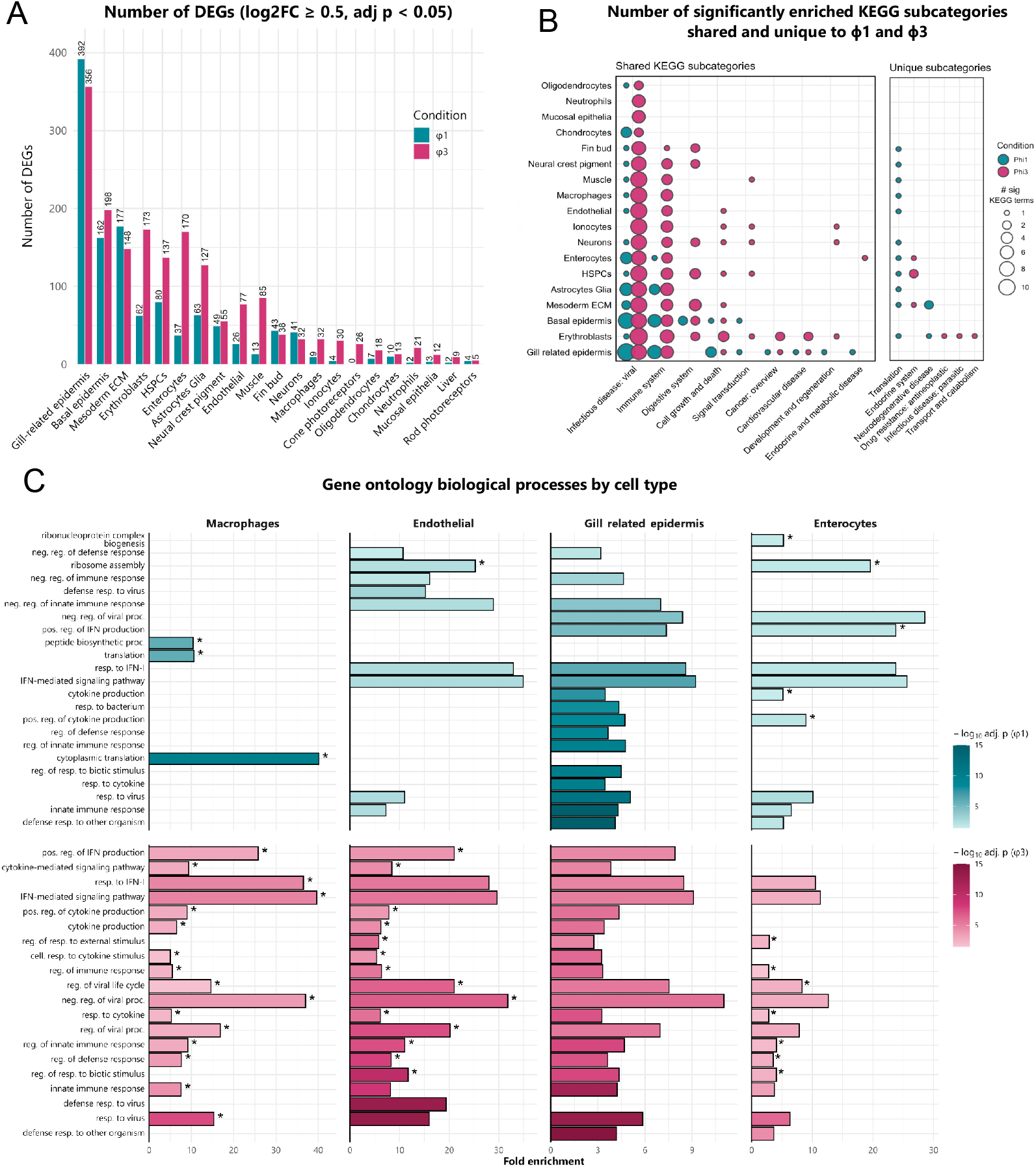
Cell-type transcriptional responses and pathway enrichment to IFNφ1 and IFNφ3. **A**. Total number of DEGs induced by IFNφ1 and IFNφ3 in each annotated cell type. DEGs were defined as genes with log2FC ≥ 0.5 and Benjamini-Hochberg FDR p adj < 0.05 **B**. Dot plot showing the number of significant enriched KEGG terms per subcategory per cell type (Benjamini-Hochberg FDR p adj < 0.05). Left: subcategories enriched in both IFNs (shared). Right: subcategories enriched in only one IFN (unique). Dot size represents the number of significant terms within that subcategory. **C**. GO Biological Process enrichment in Macrophages, Gill-related epidermis, and Endothelial cells. For each cell type and IFN group, the top 10 terms by adjusted p (after excluding predefined terms and requiring a gene count ≥ 3) define the per-row terms. All selected terms are plotted in every cell type where it passes these filters. The X axis indicates fold enrichment, defined within cell type and the fill colour represents the −log10(adjusted p) for each term as calculated by Benjamini-Hochberg FDR p adj. Asterisks mark terms uniquely significantly enriched to φ1 or φ3 treatment within that cell type. Term labels are shortened for readability and selected overlapping terms were excluded to reduce redundancy.

To understand the functional roles of either IFNφ, we used KEGG pathway analysis within each cell type. As the terms for KEGG and GO in the immune system are less well defined in the zebrafish, we generated lists of human genes orthologous to these zebrafish DEGs. This revealed cell-type specific differences in pathway induction between IFNs. To compare the distribution of enriched terms between cell types, we counted the number of significant terms in each KEGG subcategory (Figure 4B). For both IFNs, as expected the majority of terms related to viral infection and the immune system, which also represented the most enriched pathways globally (Supplementary Figure 3). IFNφ3 injection resulted in the upregulation of terms for viral infection and immune-related pathways consistently across cell types. In contrast, IFNφ1 only induced terms for both viral infection and immune responses in the gill-related and basal epidermis, Mesoderm/ECM, Astrocytes / glia, enterocytes and chondrocytes. One significant KEGG term for viral infectious disease was also present in oligodendrocytes, fin bud, neural crest pigment, Muscle, Macrophages, Neurons, HSPCs and erythroblasts. However, closer inspection of the revealed that this term was driven almost exclusively by genes related to translation, and were the same subset of genes driving activation of translation pathways in these cell types in response to φ1 (Supplementary Table 3).

For the remaining shared pathway terms, these were induced across several cell types by IFN-φ3 but restricted to the gill-related and basal epidermis for IFN-φ1. IFN-φ1 uniquely upregulated translation related terms, as well as “neurodegenerative disease” terms in mesoderm and erythroblasts, which were primarily driven by the induction of mitochondrial and ATP related genes such as ATP synthase, and cytochrome c oxidase subunits. On the other hand, IFNφ3 induced pathways related to the endocrine system in the mesoderm, HSPCs and Enterocytes, and transport and catabolism, drug resistance and parasitic infections in erythroblasts.

Interestingly, some cell types demonstrated diametrically opposing reactivity depending on the IFN subtype. Although IFNφ1 induced more terms within infectious diseases and the immune response in the gill-related epidermis than IFNφ3, these were absent or present at low levels in other cell types including cells of the immune system such as macrophages and neutrophils. The opposite pattern was present in IFNφ3 treated cells, with strong responses in macrophages and neutrophils and weaker responses in the gill-related epidermis.

Motivated by the epithelial versus immune centred specialisation of mammalian IFN-III and IFN-I (González-Navajas et al. 2012; Uccellini and García-Sastre 2018; Sommereyns et al. 2008), we examined cell types with defined barrier or systemic roles, including macrophages, enterocytes, the gill related epidermis and endothelial cells. Gene Ontology Biological Process (GO BP) enrichment on differentially expressed genes revealed that IFNφ1 primarily engages epithelial identity programs. Macrophages showed no enrichment for immune-activation terms and endothelial cells displayed only limited immune-activation with enrichment also present for negative regulation of the immune response; whereas the gill-related epidermis exhibited robust enrichment for multiple antiviral defence terms and enterocytes were enriched for the negative regulation of viral processes, as well as IFN response and cytokine production terms. In contrast, IFNφ3 elicited antiviral GO BP terms across all four cell types, suggesting that at this time point antiviral immunity is induced in a limited number of tissues in response to IFNφ1 but globally to IFNφ3.

### Evolutionarily conserved ISGs show distinct cell type expression between IFNφ1 and IFNφ3

Both the global and cell type specific KEGG pathways confirm that the primary function of both IFNφ1 and IFNφ3 is the antiviral immune response. To dissect the cell type specificities of the antiviral response at the gene level, we chose to focus on a core subset of ISGs with well-known antiviral functions, which have been shown to be evolutionarily conserved between multiple vertebrate species, including zebrafish (Levraud et al. 2019; Shaw et al. 2017).

We first selected several of these which were highly expressed in our data set and plotted these in a heatmap, which showed similar patterns of activation with all ISGs across some cell types. Enterocytes, kidney cells and other primarily epithelial tissues expressed higher levels of ISGs in response to IFNφ1, whilst mesodermal and hematopoietic tissues such as macrophages and endothelial cells consistently demonstrated increased expression to IFNφ3 (Figure 5B). Several of these demonstrated notable heterogeneity in ISG responses between IFNs, as ISGs in a given cell type did not respond uniformly. For example, the gill related epidermis had the strongest *isg15* induction in response to IFNφ1, but the opposite pattern was observed with *mxa* and *rsad2* which were expressed more strongly in response to IFNφ3 (Supplementary figure 4).

**Figure 5.**
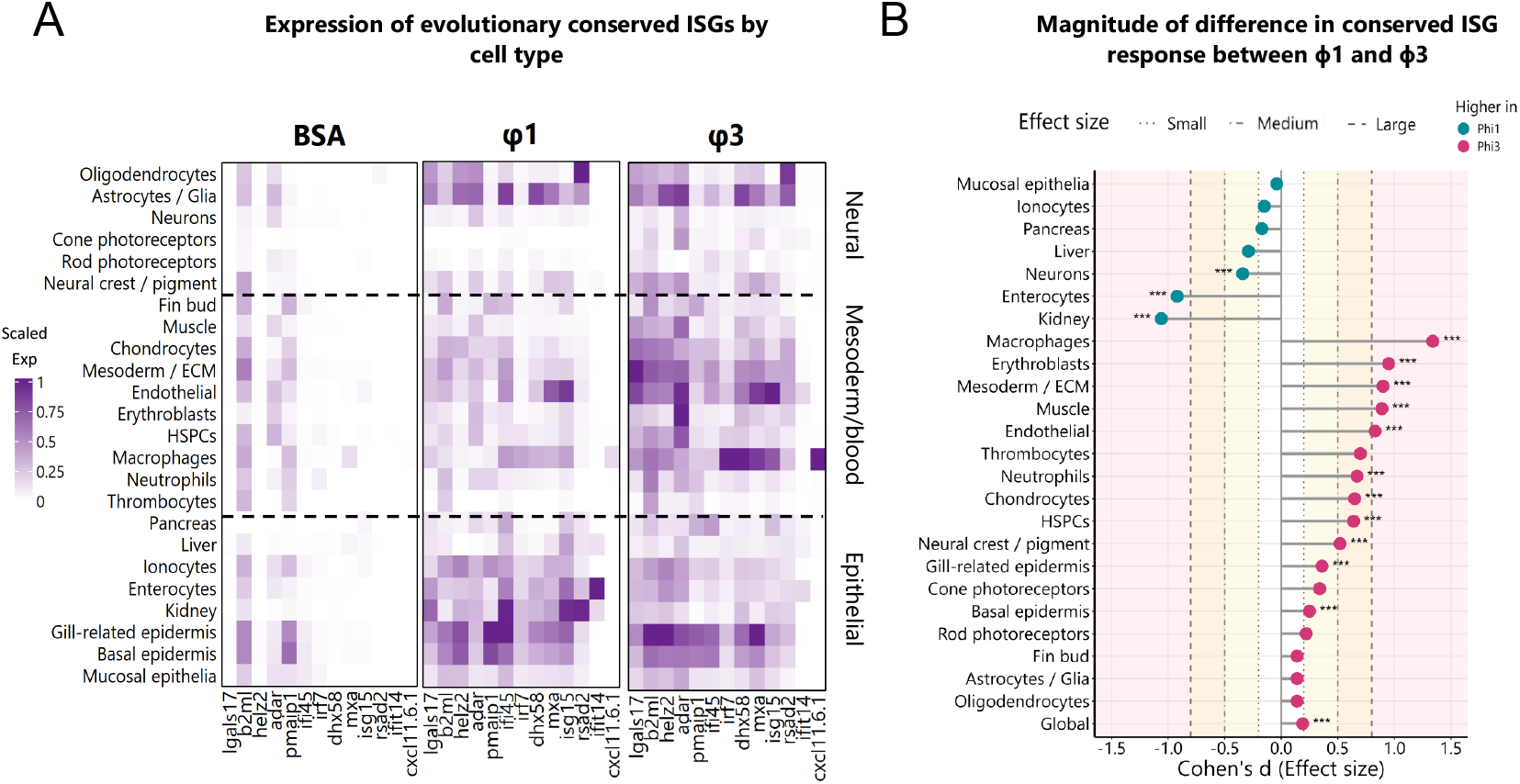
Whole larvae transcriptomics of ISG responses. **A**. Heatmap of average log-normalised expression of selected ISGs across treatment groups. Cell types are grouped broadly into neural, mesoderm/blood, and epithelial categories to highlight cell-type specific differences in ISG induction. Expression is scaled by gene to enable comparisons **B**. Lollipop plot summarizing effect size differences (Cohen’s *d*) in ISG scores between IFNφ1 and IFNφ3 for each cell type. Positive values (Magenta) indicate stronger induction with IFNφ3 and negative values (blue) indicate stronger induction with IFNφ1. Shaded background bands indicate conventional thresholds for small, medium, and large effect sizes. Asterisks mark significant differences based on one-way ANOVA followed by Tukey’s post hoc test (*p*□ <□ 0.05 *, *p*□ <□ 0.01 **, *p*□ <□ 0.001 ***).

To account for this heterogeneity when quantifying differences in the antiviral ISG response between cell types, we defined a core ISG signature based on 50 of these evolutionarily conserved ISGs, defined by their presence in Schoggins list with known zebrafish homologues that are upregulated in response to viral infection (Schoggins and Rice 2011; Levraud et al. 2019). We derived an ISG score from this signature and compared the responses to IFNφ1 and IFNφ3 using effect sizes to identify which IFN drove stronger responses in each cell type (Figure 5B). At the whole organism level, IFNφ3 resulted in a slightly but statistically significantly increased score compared to IFNφ1. Despite the overall weaker response to IFNφ1, kidney cells and enterocytes had higher ISG scores in response to IFNφ1, as well as a smaller but still significantly increased effect on neurons. Although not significant and with small effect sizes, the liver, pancreas, ionocytes and mucosal epithelia all leaned towards IFNφ1. All other cells exhibited stronger responses to IFNφ3, but this effect was strongest in immune cells, including macrophages, erythroblasts, and neutrophils, as well as mesodermal cells, muscle cells and endothelial cells. Gill-related and basal epidermis also responded more strongly to IFNφ3 with lower effect sizes.

To independently confirm the cell-type specific differences in vivo, we used double transgenic larvae expressing mCherry under the mxa promoter and GFP under the gut specific (cldn15la) or macrophage specific (mpeg1.1) promoter. Larvae were injected at 4dpf and fixed 24 hours later to account for the delay in fluorescence reporter expression noted in previous studies (Maarifi et al. 2019). Immunohistochemistry (IHC) demonstrated stronger mxa reporter expression in enterocytes exposed to IFNφ1 than to IFNφ3 (Figure 6A, B) which was confirmed by flow cytometry (Figure 6C). The opposite pattern was seen in macrophages, where IHC and flow cytometry showed stronger mxa reporter expression (Figure 6D, E) and higher numbers of mxa reporter positive macrophages (Figure 6F) to IFNφ3, indicating both an increased magnitude of ISG expression and number of responding cells.

**Figure 6.**
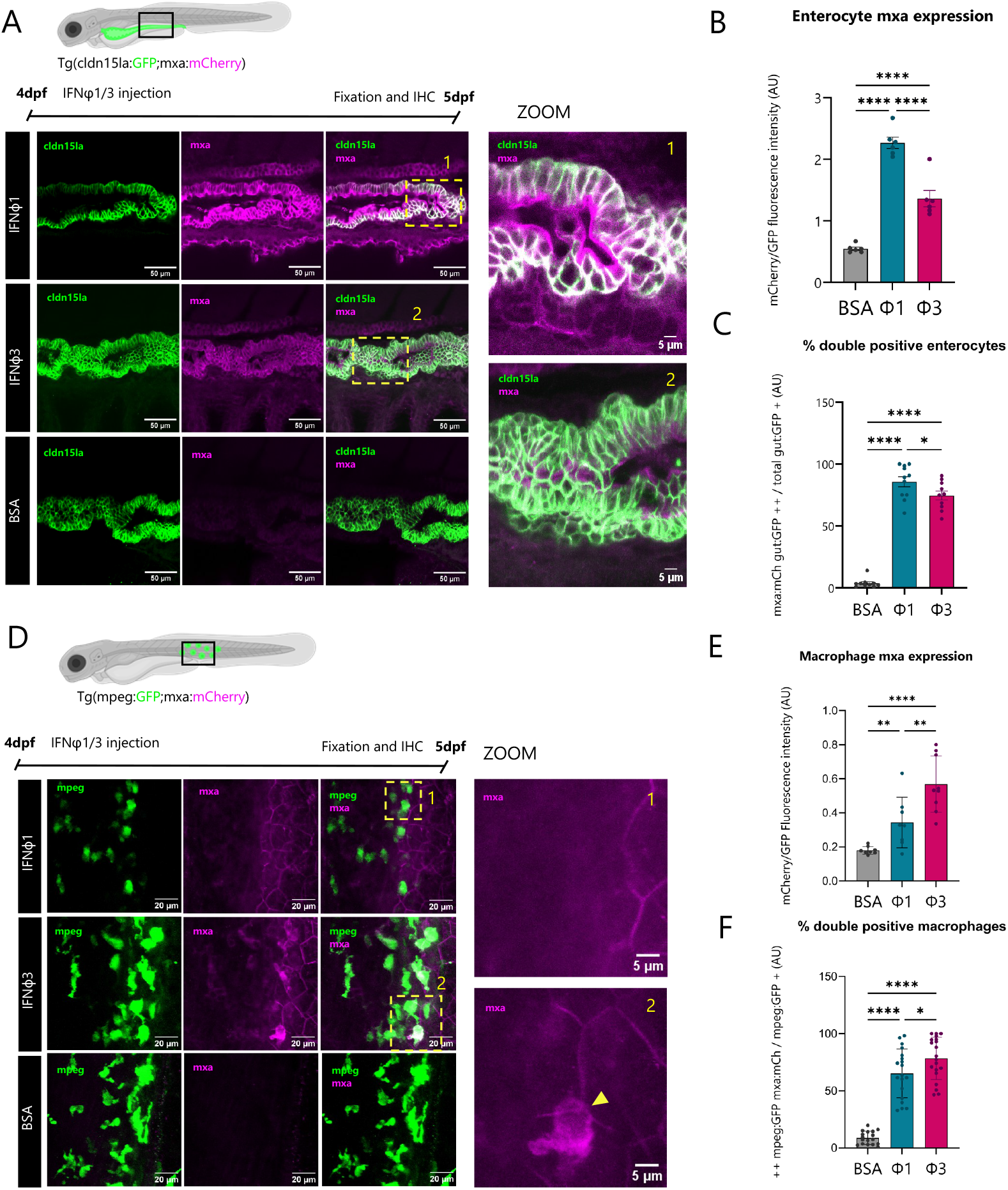
Immunohistochemistry and flow cytometry of Mxa expression in macrophages and enterocytes. **A**. Representative images of IHC against GFP in enterocytes (green) and mCherry for mxa expression (magenta) in fixed Tg(cldn5a:GFP;mxa:mCherry) larvae. Images represent a 20 slice substack. **B**. Quantification of immunofluorescence of mxa within enterocytes. Signal was quantified in 3D within a masked and segmented region of the gut and normalised to GFP signal to account for differences in cell volume/number between images (n = 6). **C**. The proportion of mxa positive enterocytes in Tg (cldn5a:GFP;mxa:mCherry) larvae as quantified by flow cytometry 24hpi following coelomic and intracerebral injection of rIFNφ1 and rIFNφ3 or control (BSA) in 4dpf zebrafish larvae (n = 7-11 replicates per condition, pools of 3 larvae per replicate). Data are scaled to concatenate 2 independent experiments. **D**. Representative images of IHC against GFP in macrophages (green) and mCherry for mxa expression (magenta) in fixed Tg(mpeg:GFP;mxa:mCherry) larvae. Images are maximum projection of z stacks. **E**. Quantification of immunofluorescence of mxa within macrophages signal was quantified in 3D within a masked and segmented region of macrophages and normalised to GFP signal to account for differences in cell volume/number between images. The dashed lines provided on Zoom images represent the outline of macrophages **F**. The proportion of mxa positive macrophages in Tg (mpeg:GFP;mxa:mCherry) larvae 24hpi as quantified by flow cytometry (n = 16-19 replicates per condition, pools of 3 larvae per replicate). Data are scaled to concatenate 2 independent experiments. All data were tested for significance using one way ANOVA with multiple comparisons. Error bars represent the SEM and asterisks indicate statistical significance *p<0.05,**p<0.01, ****p<0.0001.

### IFNφ1 and IFNφ3 are produced by different cell types during viral infection

To examine IFN induction during viral infection, we used the sindbis virus (SINV) model of encephalitis, previously shown to upregulate both IFNφ1 and IFNφ3 (Laghi et al. 2024; Boucontet et al. 2018). When injected into the pericardial cavity, SINV propagates to the central nervous system (CNS) allowing for infection both in the CNS and periphery. At 3dpf, we injected tag-BFP2 reporter SINV into double transgenic larvae producing GFP under the IFNφ3 promoter and mCherry under the IFNφ1 promoter. Imaging of the whole larvae showed that IFNφ1 was expressed in many cells throughout the larva and enriched in the caudal haematopoietic tissue, whilst IFNφ3 expression was limited to fewer cells localised showing strong expression around virus positive cells (Figure 7A). Closer examination of these cells showed that those producing IFNφ1 had similar morphology, whilst the morphology of IFNφ3 producing cells were more varied (Figure 7B). There was no evidence of co-expression of IFNφ1 and IFNφ3.

**Figure 7.**
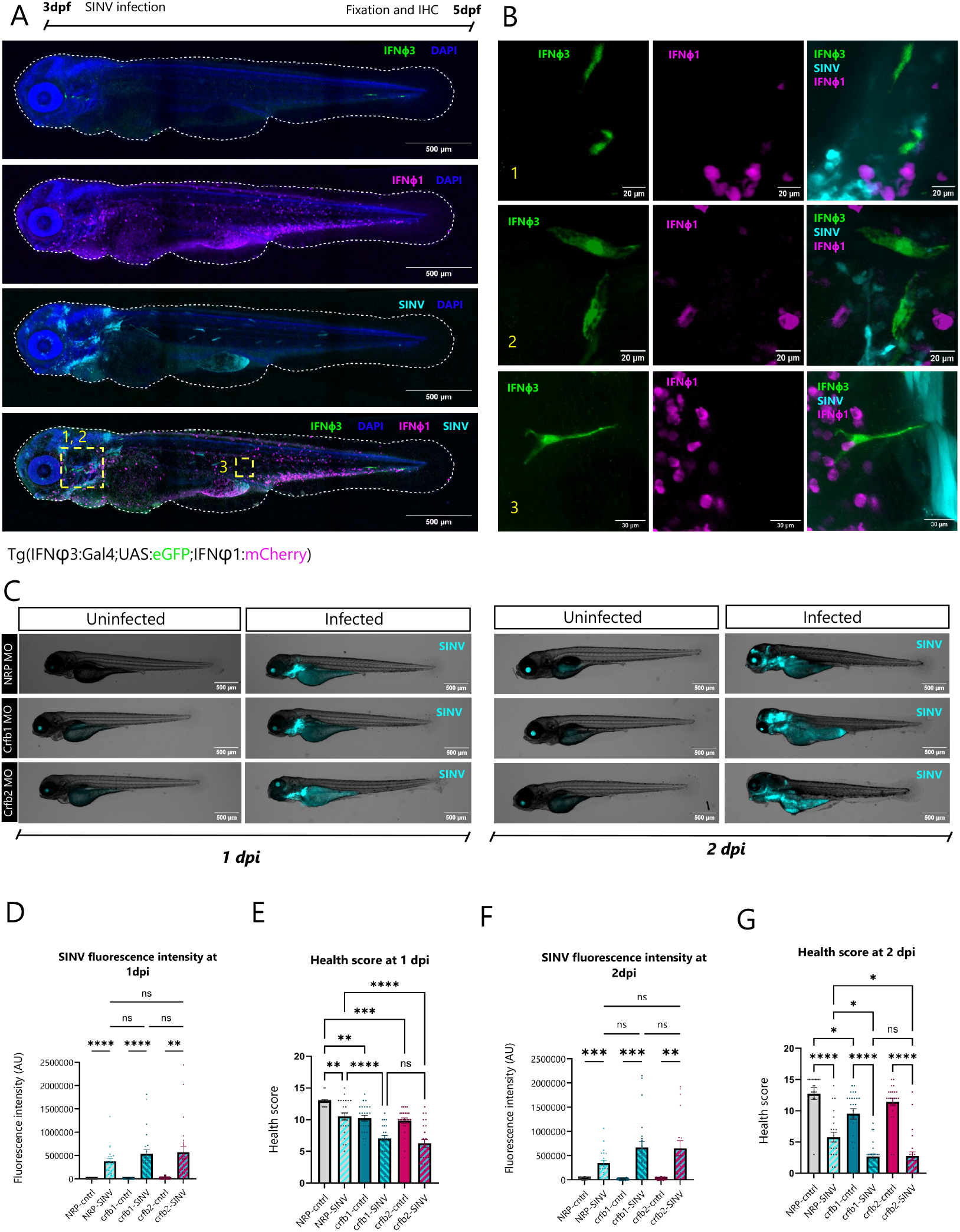
Immunohistochemistry for cell types producing IFNφ1/3 and redundancy in antiviral immunity. **A**. Representative maximum projection of z stack for fixed and IHC labelled whole Tg (IFNφ1;Gal4:UAS:GFP;IFNφ3:mCherry) larvae 2 days post infection (dpi) with SINV-TagBFP in the pericardial cavity. IFNφ1 is shown in green, IFNφ3 in magenta, SINV-TagBFP in cyan and DAPI in blue. Higher resolution imaging of individual cells expressing IFNφ1 and IFNφ3 in IHC labelled Tg (IFNφ1;Gal4:UAS:GFP;IFNφ3:mCherry) larvae. Images are maximum projections of Z stacks **C**. Widefield fluorescent and transmitted light images of crfb1, crfb2 and NRP control morphant larvae 1 and 2 dpi with SINV-TagBFP in the pericardial cavity **D**. Fluorescence intensity measurements of images of larvae 1dpi (n = 24) **E**. Health scores for larvae 1dpi (n = 24) **F**. Fluorescence intensity measurements of images of larvae 2dpi (n = 24) **G**. Health scores for larvae 2dpi (n = 24). Error bars represent the SEM and asterisks indicate statistical significance *p<0.05,**p<0.01, ****p<0.0001.

To assess if the contribution of signalling via either group conferred stronger protection against viral infection, we used morpholino knockdowns for crfb1 and crfb2, the receptors for IFNφ3 and IFNφ3 respectively. Despite a general pattern for increased fluorescence intensity at both 1 and 2pi (Figure 7C), no significant differences in fluorescence intensity (Figure D, F) or health scores (Figure E, G) were observed between the two groups and control morphant infected larvae.

## Discussion

In this work, we have identified differences in the cellular responses and antiviral programs induced in response to zebrafish IFNφ1 and IFNφ3, representative of the two subgroups of IFN-Is of teleost fish. Our results suggest that, to compensate for the loss of IFN-III in ray-finned fish, teleost IFN-Is have diversified to activate cell type specific programs paralleling those seen in mammalian IFN-I and IFN-IIIs.

Injection with IFNφ3 (group c2) drove broad and ubiquitous antiviral program across most cell types, with strongest induction in myeloid and endothelial cells, characteristic of the immune centred functions of mammalian IFN-Is (Uccellini and García-Sastre 2018; Lee et al. 2009; González-Navajas et al. 2012). The strong induction of ISGs in endothelial cells following IFNφ3 stimulation mirrors responses to IFN-I in mammals, where endothelial susceptibility to IFN-I signalling contributes directly to the pathogenesis of interferonopathies such as Aicardi-Goutières syndrome and STING-associated vasculopathy (Liu et al. 2014; Viengkhou et al. 2024). Of note, pseudobulk analysis demonstrated that chemokines, including *cxcl11*, were only induced globally by IFNφ3, consistent with data in mammalian models that demonstrate CXCl10 and CXCL11 regulation in response to IFN*β* but not IFN-III (Forero et al. 2019).

Functional enrichment demonstrated that IFNφ3 drove pathways for the antiviral immune response across all cell types, whilst simultaneously inducing diverse pathways including cell growth and death, as well as digestive and endocrine system pathways, which were largely absent in response to IFNφ1. This is consistent with clinical data that shows that recombinant IFN*α* but not IFN-III induces high endocrine dysfunction and thyroiditis (Tomer and Menconi 2009; Fredlund et al. 2015).

In contrast IFNφ1 mediated responses were restricted predominantly to epithelial tissues, including the gut, kidney, and various epithelial subsets, where canonical ISGs and immune pathways were activated. These are prototypical IFN-III responsive tissues, consisting primarily of epithelial cells and is consistent with literature demonstrating strong ISG expression in the tubular kidney and intestinal epithelia in response to IFN-III (Sommereyns et al. 2008; Mahlakõiv et al. 2015; Hermant et al. 2014; Goel et al. 2020).

Alongside the cell type restriction, the transcriptional response triggered by IFNφ1 was comparatively limited in scope, and constituted largely of a subset of the genes induced by IFNφ3, resembling the restricted transcriptional profile observed for IFN-III relative to IFN*α* in mammals (Zhou et al. 2007). Notably, the 3 genes uniquely upregulated by in response to IFNφ1 have all been linked to epithelial tissues. Ctrb1 is digestive serine protease expressed in the exocrine pancreas, a ductal epithelial tissue (Zhang et al. 2014), nppa encodes atrial natriuretic peptide, receptors for which have been demonstrated on epithelial tissues involved in osmoregulation such as the branchial arches and pronephric ducts (Gong et al. 2018). Finally, *namptb* driven inflammation increases keratinocyte levels of reactive oxygen species to drive epidermal pathology in zebrafish models of inflammatory skin diseases, which can be rescued by nampt inhibition (Martínez-Morcillo et al. 2021).

The magnitude of canonical ISG expression to IFNφ1 and IFNφ3 in these tissues largely reflected this “barrier or not barrier” role, but was not absolute. Whilst prototypical IFN-III responsive tissues, such as the gut and kidney showed a clear preference for IFNφ1, other epithelial compartments such as basal and gill-associated epidermis exhibited stronger induction by IFNφ3, though with smaller effect sizes. This may be explained by the immaturity of gills compared to other barrier organs at the larval stage we studied (Kämmer et al. 2022). Viral restriction in the gut is heavily dependent on IFN-III but not IFN-I responses (Mahlakõiv et al. 2015), whereas keratinised epithelia has been shown to be affected by both IFN-I and IFN-IIIs, with humans even having a keratinocyte specific IFN-I, termed IFN*κ* (LaFleur et al. 2001). Accordingly, active roles for both IFN-I and IFN-III at keratinised epithelia have been demonstrated during health and disease (Sarkar et al. 2018; Bernard et al. 2015; Goel et al. 2020; Zahn et al. 2011).

Additionally, we observed IFNφ1–dependent induction of select ISGs in non-epithelial cell subsets. Although whole-organism comparisons of IFN-I versus IFN-III remain limited, available data support a degree of non-epithelial responsiveness. IFN-III elevates ISG expression in neutrophils (Stifter et al. 2019) and plasmacytoid dendritic cells (Kelly et al. 2016). The neuronal IFN-III response has been found to be protective against herpes viruses (Salazar et al. 2023) and primary human neurons and astrocytes in culture respond to recombinant IFN-III in an il10rb dependent manner, consistent with direct responsiveness (Li et al. 2010). However, in the olfactory system, epithelial cells but not olfactory neurons responded to IFN-III suggesting that neuronal responsiveness is restricted to certain neuron subsets (Jacobs et al. 2019).

In mammals, differences in transcriptional breadth have been suggested to reflect altered magnitude and kinetics of the response triggered by IFN types, where universal IFN-Is dominate early systemic responses and IFN-IIIs provide slower, persistent signalling at epithelial barriers (Pervolaraki et al. 2018; Forero et al. 2019; Dickensheets et al. 2013). Kinetically, our data diverge from the mammalian model, as the epithelial biased IFNφ1 resulted in shorter lived ISG expression compared with IFNφ3. Interestingly, studies in adult zebrafish report more mammal-like kinetics, with IFNφ3 inducing rapid and transient responses and IFNφ3 driving slower transcriptional programs (López-Muñoz, F. J. Roca, et al. 2009). This suggests other factors may play a role in ISG kinetics, such as developmental stage, ISG readout choice or even the presence of adaptive immunity in adults that is absent in larvae.

Our data further indicate that IFNφ1 and IFNφ3 are produced by distinct cellular sources during viral infection. In the SINV model, IFNφ1 was broadly expressed in cells enriched within the caudal haematopoietic tissue, suggesting a myeloid origin. This is consistent with previous work identifying neutrophils as the major source of IFNφ1 during infection with another alphavirus, Chikungunya virus (Palha et al. 2013). In contrast, we observed IFNφ3 expression in cells of varied morphology. The exact identity of IFNφ3-producing cells is unknown, although we observed one IFNφ3-expressing cell consistent with neuronal morphology, suggesting IFNφ3 may be produced by diverse or rare cell types. We also saw no evidence of co-expression of IFNφ1 and IFNφ3, suggesting that they are regulated independently. In response to spring viremia carp virus, it has been shown that IFNφ1 and IFNφ3 are regulated by different IRF dependencies. Cytosolic PRRs pathways favour IRF3 recruitment to IFNφ1 promoters and TLR/MyD88 pathways favour IRF7/1 complex formation at IFNφ3 promoters (Feng et al. 2016). Defining these sources will be critical for understanding the spatial and temporal dynamics of antiviral IFN responses in zebrafish. Distinct cellular sources for different teleost fish IFNs have been observed in the much more complex situation of salmonids (Svingerud et al. 2012).

One limitation of this work is the lack of mechanistic insight into the cell type specificity. In mammals, this is heavily dependent on differential receptor localisation (Sommereyns et al. 2008). Unfortunately, we’ve been unable to resolve receptor localisation due to very low transcript abundance, either by in situ hybridization or by interrogating available single cell RNA-based atlases. Beyond receptor expression, other mechanisms likely affect the cell type differences between IFNφ1 and IFNφ3. Epigenetic mechanisms, as well as cell polarisation have been found to influence the relative responsiveness of intestinal epithelial cells to IFN-III and IFN-Is in mammals (Bhushal et al. 2017).

Finally, a remarkable outcome of this study was not only the overall differences in responses to subtypes, but also the strong heterogeneity in ISG expression, which did not always follow consistent patterns. The fact that the ISG repertoire is strongly cell type-specific may provide insight into diseases with tissue selective vulnerability despite widespread receptor expression, including type I interferonopathies which in humans primarily affect the skin and CNS (Crow and Stetson 2022). Identification of cell type specific ISGs in these tissues, as well as the mechanisms for cell type specificity will be essential for understanding pathological outcomes and translational applications.

Our findings suggest that the two main teleost type I IFN groups, represented by zebrafish IFNφ1 and IFNφ3, have independently acquired cell type specialisation seen between mammalian IFN-I and IFN-IIIs. Furthermore, IFN subtypes are produced by distinct subsets of cells, suggesting that the division of labour between IFN groups is defined by both cellular responsiveness and the identity of producing cells. The independent emergence of epithelial and immune centred IFN programs suggests that cellular separation of antiviral immunity is highly evolutionarily advantageous, providing strong selective pressure for its convergent evolution in different vertebrate lineages. Our work also has implications in the use of zebrafish as model organisms in IFN-driven pathologies, highlighting the importance of resolving functional divisions of paralogues in zebrafish.

## Materials and methods

### Zebrafish lines and husbandry

All experiments were conducted according to the European union directives for animal research and were approved by the Ethics Committee of Institut des Neurosciences Paris-Saclay (CEEA n°59) under approval #39238. The genetic background of all fish used was AB carrying one or two copies of the nacre mutation (*mitfa*^*w2*^ allele), obtained by at least 7 backcrosses of nacre mutants onto AB. This resulted in fish either with a wild-type striped colour pattern (w2/+) or without body stripes (w2/w2). Eggs were obtained by natural spawning Typically; striped males were crossed with nacre females or vice-versa; the colour difference allowed easy re-identification of males and females post-mating.

A new *zps2* allele of the Tg(*isg15:eGFP; myl7:eGFP*) ISG15 reporter line (Balla 2020) was generated by Tol2 transgenesis into AB eggs. Other transgenic lines used included the MXA reporter Tg*(cryaa:DsRed,mxa:mCherryF)*^*ump7*^ (Maarifi et al, 2019), the enterocyte reporter TgBAC(*cldn15la:eGFP*)^*pd1034*^, the macrophage reporter Tg(*mpeg1:EGFP*)^*gl22*^ and the IFNφ1 reporter *Tg(ifnphi1:mCherry)*^*ip1*^*(Palha 2013)*. Following collection, eggs were sterilised by treatment with 0.003% bleach and rinsed with embryo medium (E3). For the first 24 hours eggs are raised in E3 supplemented with 0.01% methylene blue. After 24 hours, the medium was changed to standard E3 without methylene blue or for experiments with live imaging, E3 with 1X PTU ((1-Phenyl 2-Thiourea: P7629; Sigma-Aldrich) to prevent pigment formation.

### Recombinant IFNs

Recombinant IFNs were produced in *Escherichia coli* as described previously (Aggad 2009), as soluble proteins in PBS pH7.4 plus 10% glycerol; the final IFNφ1 stock was at 2.5mg/ml while IFNφ3 was at 0.8mg/ml. The main IFN stocks were stocked at −80°C while one working aliquot was kept at −20°C.

### Virus production

The SINV-TagBFP2 virus was generated onto the same TE12 backbone used for the SINV-GFP strain generated previously (Boucontet et al. 2018). The tagBFP2 sequence (Subach et al. 2011) was synthetized by vector builder and cloned into the SINV genomic cDNA after a TE2 self-cleaving peptide, within the ORF coding form viral structural proteins between capsid- and envelope-coding sequences. Viral mRNA was generated by in vitro transcription from this plasmid using a SP6 mMessage mMachine kit (Thermo Fisher). This synthetic mRNA was injected into 1-cell stage AB zebrafish zygotes. After 24 hours of development at 28°C, 20 embryos with strong blue fluorescence, all with abnormal development, were dissociated into 100µL of PBS 1X, and the lysate was cleared by a brief centrifugation and filtered onto a 0.2µM syringe-held filter. 25µL of this filtrate was used to seed a confluent monolayer of BHK-21 cells (ATCC CCL-10). After 48 h at 33°C, when cells started to detach, the supernatant was collected and cleared by 10 minutes of centrifugation at 2000g, aliquoted and frozen, to generate the SINV-TagBFP2 stock, at an approximate titer of 3.10^7^ PFU/mL.

### Microinjection

Larvae were anaesthetised using Tricaine at 240 µg/mL (MS222 A5040; Sigma-Aldrich). GC100F-10 capillaries (Harvard Apparatus) were pulled (heat 535, pull 30, velocity 120, time 200) using a glass micro-pipette puller (Sutter Instrument, Model-P97). Capillary tips were manually broken and airflow adjusted to achieve desired injection volume. For injection of recombinant IFNs 4dpf larvae were positioned on an agarose plate and injected with 1nl of either 1 mg/ml rIFNφ1, or 0.8 mg/ml rIFNφ3 or 1 mg/ml BSA into the coelomic cavity and hindbrain. For SINV infections larvae were injected with 2nl (∼60 PFU) SINV-TagBFP2 into the pericardial cavity at 3dpf.

### Morpholino knockdowns

Morpholinos have been previously described (Levraud et al. 2007). 1nl (4ng) of morpholino antisense oligonucleotides (Gene Tools, Philomath, OR, USA) was injected into the cell or yolk of AB embryos at the 1 to 2 cell stage. Crfb1 splice morpholino (CGCCAAGATCATACCTGTAAAGTAA) was used to knock down the group 1 (IFNφ1) IFN receptor, and crfb2 splice morpholino (CTATGAA TCCTCACCTAGGGTAAAC) to knock down the group 2 (IFNφ3) IFN receptor. Control morphants were injected with 4ng control morpholino, with no known target (GAAAGCATGGCATCTGGAT CATCGA).

### Immunohistochemistry

Larvae were euthanised by anaesthetic overdose (tricaine 800µg/mL for 5 minutes) and fixed overnight in 1X PBS, 4% PFA, 0.1% Triton, 0.05% sodium azide 4 °C. Immunostaining was performed as described previously (Lempereur et al. 2022), with a modified depigmentation solution containing 1X PBS, 5% H202, 0.1% Triton, 0.05% sodium azide. Staining’s were performed against fluorophores (GFP for eGFP, dsRed for mCherry) for transgenics and TagFP for SINV, primary antibodies (chicken anti-GFP Aves labs GFP-1020 1:300, rabbit anti-dsRed clontech Takara 632496 1:300, anti-TagFP 647 Synaptic systems N0502-At488-L 1:300) and secondary antibodies (anti-chicken alexafluor 488 invitrogen A11039 1:500, anti-rabbit alexa 546 invitrogen A11010 1:500). DAPI (1:1000) was added alongside secondary antibodies.

### qPCR

Larvae were euthanised by anaesthetic overdose and pooled into groups of 3 prior to lysis at 1, 2, 4, 6, 8, 24 and 48 hours. RNA was extracted by addition of 300 μL of RLT buffer (Qiagen) and mechanical dissociation by pipetting. Total RNA was extracted using the Qiagen RNA easy mini kit following the manufacturer’s instructions and was eluted in nuclease free water. Reverse transcription was performed on 10 μL of eluted RNA using the SuperScript IV Reverse Transcriptase kit with dT17 primer. cDNA was diluted to a final volume of 200 μL (10x), of which 5 μL was used as a template for each qPCR assay using the PowerUp SYBR Green master mix for quantification. (ThermoFisher Scientific) with primer pairs (Table 1). The QuantStudio™ 5 System was used for real-time qPCR with the following cycling parameters: 50°C for 2 mins, 95°C for 10 mins, 40 cycles of 95°C for 15s and 60°C for 1 minute. Results are reported as relative transcript expression compared to a reference cDNA with normalisation performed using the housekeeping gene elongation factor 1 alpha 1 (ef1a). The mitochondrial gene MTCB was used to assess sample quality and samples with high levels were excluded from downstream analysis.

### Low magnification imaging of whole larvae

Larvae were anaesthetised using 1.5X Tricaine in E3 and mounted laterally or dorsally on moulds made of 1% Agarose in E3. Imaging was performed on an EVOS widefield microscope (Thermo Fisher Scientific) using a 2× objective with transmitted light and TxRed/GFP fluorescence filter sets.

### Confocal imaging

Fixed larvae were mounted laterally or dorsally on moulds made of 1% Agarose in E3. and imaged on a Leica SP8 confocal microscope (HC FLUOTAR L 25×/0.95 water-immersion and HC PL FLUOTAR 10X/0.30 dry objectives). Excitation was provided by 405nm, 488 nm, 552 nm and 638nm and lasers. For imaging of mxa in enterocytes Tg(cldn15la:GFP;mxa:mCherry), the 25X objective at a resolution of 1024×1024 with a frame average of 2 and system optimised Z step of 0.684μm with 488nm and. For mxa in macrophages Tg(mpeg1.1:GFP;mxa:mCherry) the 25X objective at a resolution of 1024×1024 with a Z step of 1μm, 3X optical zoom and excitation by 488nm and 638nm lasers. For whole Tg(IFNphi3:UAS;Gal4:GFP;IFNphi1:mCherry) the 10X objective with a 0.75× digital zoom. Image stacks were acquired at a resolution of 1024 × 1024 with a z-step size of 5.0 µm. Excitation was provided by 405 nm, 488 nm, 552 nm, and 638 nm lasers. For higher resolution Tg(IFNphi3:UAS;Gal4:GFP;IFNphi1:mCherry) the 25X objective with a 3× digital zoom. stacks were acquired at a resolution of 1024 × 1024 with a z-step size of ∼0.5 µm. Excitation was provided by 405 nm, 488 nm, 552 nm, and 638 nm lasers.

### Image analysis

Image analysis was performed on FIJI. Fluorescence intensity was quantified in macrophages and enterocytes by thresholding to generate binary masks. Thresholding and mask generation were standardised and automated using experiment-specific macros to ensure consistency between images. The Fiji Image calculator was used to multiply binary masks by fluorescence channels to restrict signal measurements to masked cells. Fluorescence intensities were normalised by calculating the ratio of the signal of interest relative to the reporter signal to account for variability in cell number/volume.

For quantification of fluorescence intensity in SINV infected fish, thresholding and fluorescence intensity quantifications were performed by day using standardised settings on quantifish (Stirling 2020).

### Flow cytometry

Larvae were over anaesthetised in 5X tricaine for 5 minutes. Whole larvae were dissociated using a combination of enzymatic and mechanical dissociation, using 1X trypsin and the gentleMACs tissue dissociator (Miltenyi Biotech) in a C tube for 40seconds at an rpm of 624. Following dissociation cells were kept at room temperature for 5 minutes before addition of 5ml of cold 1XPBS with 0.04% BSA 1% FBS. The cell solution was then centrifuged for 5 minutes at 300 rcf before resuspension in 500ul of cold 1XPBS with 0.045 BSA and 1%FBS. The solution was passed through a 40um cell strainer (ref) to discard large debris and 1X Draq7 (Thermo Fisher Scientific D15106) was added to the cells to label dead cells. Flow cytometry was performed using the Attune™ NxT Acoustic Focusing Cytometer (Thermo Fisher Scientific). Data were analysed using FlowJo™ Software (v10.x). Doublets and debris were excluded based on FSC-A vs. FSC-H and SSC, dead cells removed on the basis of Draq7 positivity. Pigment cells were removed on the basis of autofluorescence in non-fluorescent controls and endogenous fluorescence of reporter lines was used to identify cell types and gating thresholds were set based on wild type or unstained controls, and applied uniformly across all samples, see supplementary figure 5 for gating strategy.

### Health scoring

Health scoring was performed as previously described (Palha et al. 2013), clinical signs of infection including equilibrium, response to touch, body shape, blood flow, heartbeat, oedema, swim bladder presence and yolk opacity were scored in infected animals.

### Single cell isolation and library preparation

Larvae were over anaesthetised in 5X tricaine for 5 minutes. Whole larvae were dissociated using a combination of enzymatic and mechanical dissociation, using the gentleMACs tissue dissociator (Miltenyi Biotech) with 1 ml of dissociation solution (Buffer X 0.95X, Enzyme P 0.025X, Buffer Y 0.010X, Enzyme A 0.005X) prepared from the Neural P dissociation kit (Miltenyi Biotech). Following dissociation cells were kept at room temperature for 5 minutes before addition of 5ml of cold 1XPBS with 0.04% BSA 1% FBS. The cell solution was then centrifuged for 5 minutes at 300 rcf before resuspension in 500ul of cold 1XPBS with 0.045 BSA and 1%FBS. The solution was passed through a 40um cell strainer to discard large debris. Dead cells were removed by magnetic separation using the octomacs magnet and Dead Cell Removal Kit (Miltenyi Biotech) according to the manufacturer’s instructions. The unlabelled flow through after magnetic separation was collected as the viable fraction and cells were resuspended into 1XPBS with 0.04% BSA checked for viability using propidium iodide.

Chromium Single Cell 3’ libraries were prepared on the Chromium iX machine and processed according to the manufacturer’s instructions to capture 10,000 cells per sample. Sequencing was performed using the NextSeq2000 P4 cell. Sequencing data has been deposited onto ArrayExpress under accession number E-MTAB-15833.

### Bioinformatics

Raw sequencing data were demultiplexed using bcl-convert and adaptor sequences were trimmed with cutadapt. Read quality was assessed with FastQC and aggregation of 2 independent experiments, gene alignments and transcript quantification were done using cell ranger. Bioinformatic analysis was performed in R studio using Seurat (V.5.2.1) and cluster profiler. Doublets and cells with >15% mitochondrial gene content were filtered out to remove low quality cells. For umap generation PCA and dimensionality reduction was performed before graph-based clustering using the top 25 PCs at a resolution of 0.5. Clusters were manually annotated according to marker gene expression and the Farnsworth atlas for the developing zebrafish embryo (Farnsworth et al. 2020) genes used for identification are highlighted in supplementary data table 2 and their expression in supplementary figure 1. Clusters enriched for RBC markers were identified based on high expression of *hbbe1*.*2* and low *mki67* and removed before re-clustering the dataset using the top 25 PCs and a resolution of 0.6.

#### Pseudobulk analysis

Raw counts were aggregated per sample to generate pseudobulk profiles. Differential expression between treatment groups and control (BSA) was modelled with DESeq2 using a group design. Genes with total counts ≤10 across samples were removed prior to fitting. Effect sizes were reported as log2 fold changes shrunken using adaptive shrinkage (ashr) (Stephens 2017). P values were adjusted for multiple testing using the Benjamini Hochberg false discovery rate (FDR).

#### Cell type DEG analysis

For each annotated cell type, treatment groups (Phi1, Phi3) were compared against the control (BSA) using the MAST hurdle model. Groups were randomly downsampled without replacement to the size of the smallest group within each contrast. Comparisons with fewer than 10 cells per group were not tested and batch was included as a covariate in DEG analysis to account for interexperiment differences. Genes were tested if detected in ≥10% of cells in either group, with no log fold-change threshold. P values were adjusted for multiple testing using the Bonferroni method.

#### KEGG and GO

For orthology based GO and KEGG pathway analysis, Zebrafish genes from differential expression result tables were mapped to human NCBI Gene IDs to enable KEGG and GO enrichment using human annotation databases. Human KEGG and GO pathway annotations were used for the mapped genes to account for limited annotation of zebrafish immune pathways. Initial orthologue assignment was performed in Python using the danRerlib package(Schwartz et al. 2024). Genes not assigned an orthologue by this method were manually checked against the ZFIN database. For genes with a reported human orthologue in ZFIN, the corresponding human NCBI Gene ID was added to a manually curated mapping file. This mapping file was then used to map all remaining DEG result tables in R to fill missing orthologues. For genes with multiple predicted human orthologues, the first orthologue listed in ZFIN was selected. This approach has limitations, as zebrafish often possess multiple paralogues, which may result in duplicated human gene assignments. To account for this only unique Human NCBI gene IDs were used for KEGG and GO pathway analysis to prevent statistical conflation. DEGs were filtered to retain upregulated genes with adjusted P ≤ 0.05 and log2FC ≥ 0.5 and cell types with fewer than 10 DEGs were excluded from enrichment analysis. Enrichment was performed using over representation analysis for Gene Ontology Biological Process and KEGG human pathways, using Human NCBI Gene IDs. The background was defined per cell type as all genes expressed in that cell type (≥5% detection). P values were adjusted for multiple testing using the Benjamini Hochberg false discovery rate (FDR). For plotting only terms with P adj < 0.05 and count ≥ 3 were used.

#### ISG module scoring and statistical analysis

A list of 50 evolutionary conserved ISGs defined by their presence in Schoggins list (Schoggins and Rice 2011) and in (Levraud et al. 2019) and was used to compute a per-cell ISG score using Seurat’s module score function. Differences in score between groups were assessed by one-way ANOVA, followed by Tukey’s HSD test for pairwise comparisons. Analyses were performed globally across all treatment groups and within each cell type. Effect sizes for pairwise comparisons were summarised with Cohen’s d (pooled SD), and plotted with directionality.

## Supplementary figure legends

**Supplementary figure 1. Cluster markers used for annotation**

Heatmap shows average expression of predefined marker genes within annotated cell types. Marker genes were grouped by cell type and filtered to those present in the object. For each gene, average expression was computed using log-normalised expression for all cells in each cell type and scaled across all cell types.

**Supplementary figure 2. Umap split by treatment group**

UMAP projection of cells from each treatment group across the clusters obtained after QCs and removal of erythrocyte clusters and doublets. Dimensional reduction was performed using PCA on the top 2,000 variable genes (25 PCs). Clustering was performed using the Louvain algorithm at resolution 0.6.

**Supplementary figure 3. KEGG and GO pathway analysis of global responses amongst pseudobulk DEGs**

A. KEGG pathways enriched in the whole larvae in response to IFNφ1 and 3. The axis shows the gene ratio, dot size represents the gene count and the fill colour represents the −log10(adjusted p) for each term as calculated by Benjamini-Hochberg FDR p adj. B. GO Biological Process enrichment in in whole larvae. For each cell type and IFN group, significant terms (after excluding predefined terms and requiring a gene count ≥ 3) define the per-row terms. All selected terms are plotted in every cell type where it passes these filters. The X axis indicates gene count and the fill colour represents the −log10(adjusted p) for each term as calculated by Benjamini-Hochberg FDR p adj. Terms that are unique to one IFN group are highlighted in bold.

**Supplementary figure 4 – Violin plots of cell type expression of rsad2, isg15 and mxa**

Violin plots of log-normalized expression of *isg15, mxa*, and *rsad2* across annotated cell types for each treatment condition. Violins show distribution of expression within each cell type and the central dot represents the mean expression

**Supplementary figure 5 – Gating strategy for the identification of mxa positive cells / enterocytes and macrophages**

A. Cells identified using FSC-A/SSC-A B. Singlets defined by FSC-H/FSC-A C. Live cells were defined based on gating of samples with and without the dead cell strain Draaq7 RL2-A/FSC-A. D. Autofluorescence from pigment cells removed by gating of pigmented non-fluorescent samples compared with GFP positive samples. E. Quadrants for GFP and mCherry positive cells were defined according to non-fluorescent controls.

## Acknowledgements

We would like to thank the zootechnicians of TEFOR Paris-Saclay for the care of the fish and Dorian Champelovier for assistance in microscopy experiments. We are also indebted to Laure Riffault from the NeuroPSI transcriptomics platform for library preparation, and Yan Jaszczyszyn and Delphine Naquin at I2BC for the sequencing and initial bioinformatic analysis. The Tg(isg15:EGFP; myl7:EGFP) construct was kindly provided by Dr Nels Elde (University of Utah). Recombinant zebrafish interferons have been produced by the recombinant Protein Production and Purification platform (PFP3R) of Institut Pasteur, Paris.

## Author contributions

Conceptualization: HW, JPL

Data Curation: HW, JPL

Formal Analysis: HW, DP

Funding Acquisition: JPL, GL

Investigation: HW, DP, VB, TB, MAY, IC, GL, JPL

Methodology: HW, GL, JPL

Project Administration: JPL

Resources: VB, TB, MAY, GL, JPL

Software: HW

Supervision: JPL

Validation: HW, JPL

Visualization: HW, DP

Writing – Original Draft Preparation: HW

Writing – Review and Editing: DP, JPL (for the moment)

