## Supplementary Figures 1-4 for "The two groups of zebrafish type I interferons target different tissues, paralleling the mammalian type I: type III IFN functional division"

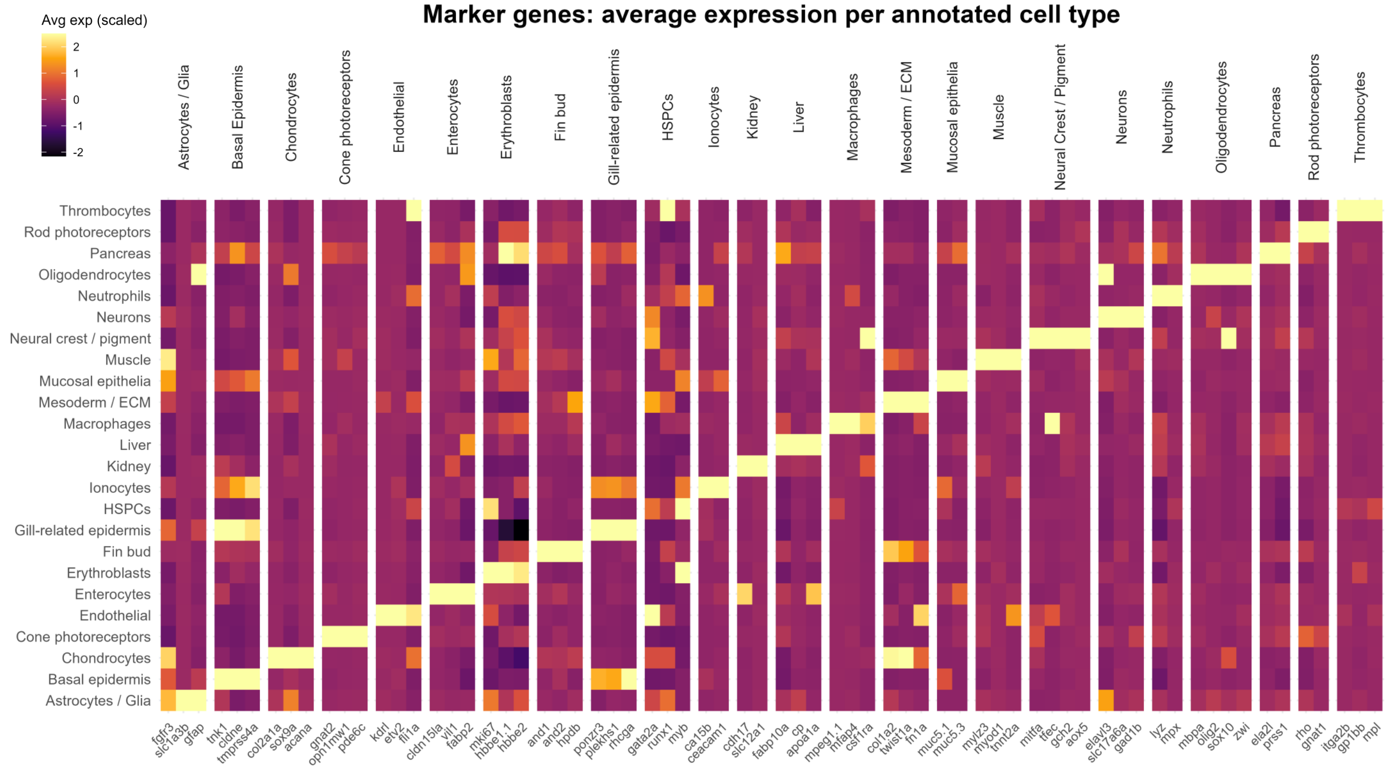


### Supplementary figure 1. Cluster markers used for annotation

Heatmap shows average expression of predefined marker genes within annotated cell types. Marker genes were grouped by cell type and filtered to those present in the object. For each gene, average expression was computed using log-normalised expression for all cells in each cell type and scaled across all cell types.

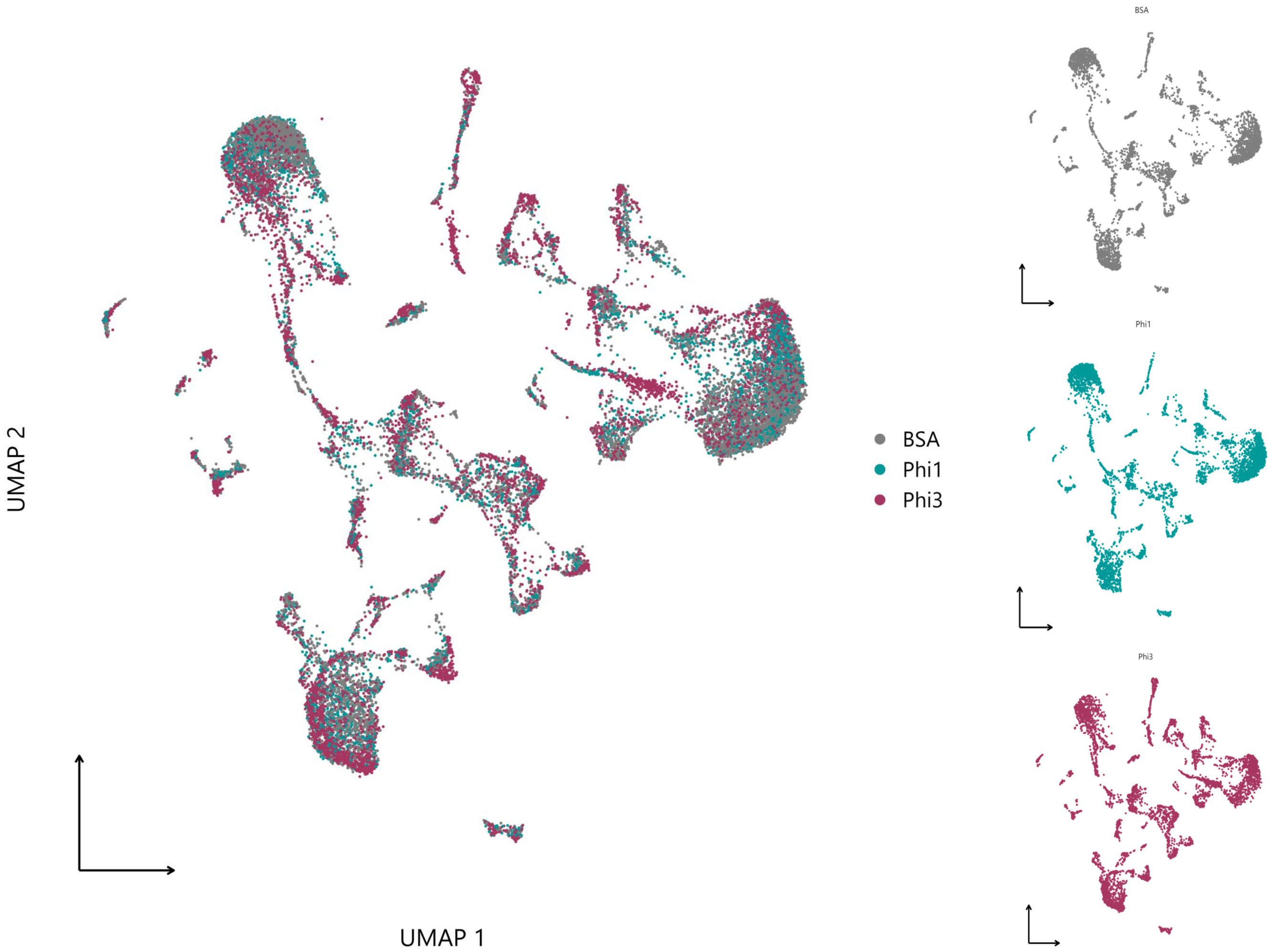


**Supplementary figure 2. Umap split by treatment group**

UMAP projection of cells from each treatment group across the clusters obtained after QCs and removal of erythrocyte clusters and doublets. Dimensional reduction was performed using PCA on the top 2,000 variable genes (25 PCs). Clustering was performed using the Louvain algorithm at resolution 0.6.

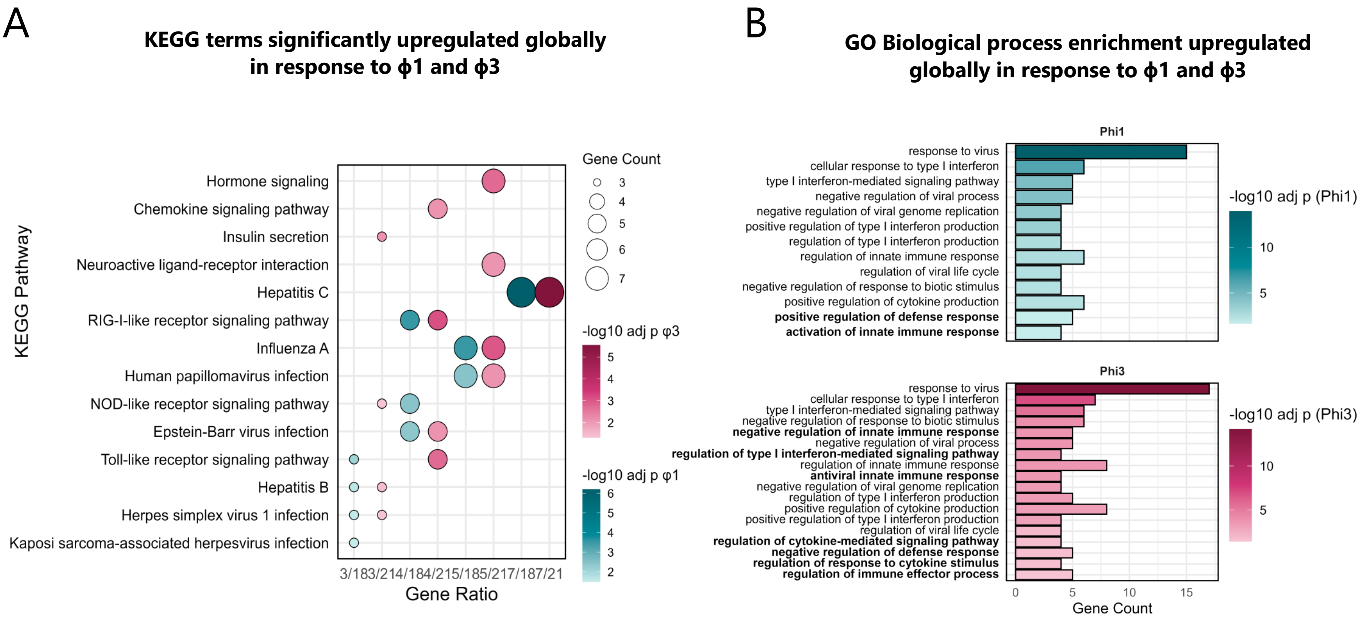


**Supplementary figure 3. KEGG and GO pathway analysis of global responses amongst pseudo-bulk DEGs**

**A.** KEGG pathways enriched in the whole larvae in response to IFNφ1 and 3. The axis shows the gene ratio, dot size represents the gene count and the fill colour represents the −log10(adjusted p) for each term as calculated by Benjamini-Hochberg FDR p adj. **B.** GO Biological Process enrichment in in whole larvae. For each cell type and IFN group, significant terms (after excluding predefined terms and requiring a gene count ≥ 3) define the per-row terms. All selected terms are plotted in every cell type where it passes these filters. The X axis indicates gene count and the fill colour represents the −log10 (adjusted p) for each term as calculated by Benjamini-Hochberg FDR p adj. Terms that are unique to one IFN group are highlighted in bold.

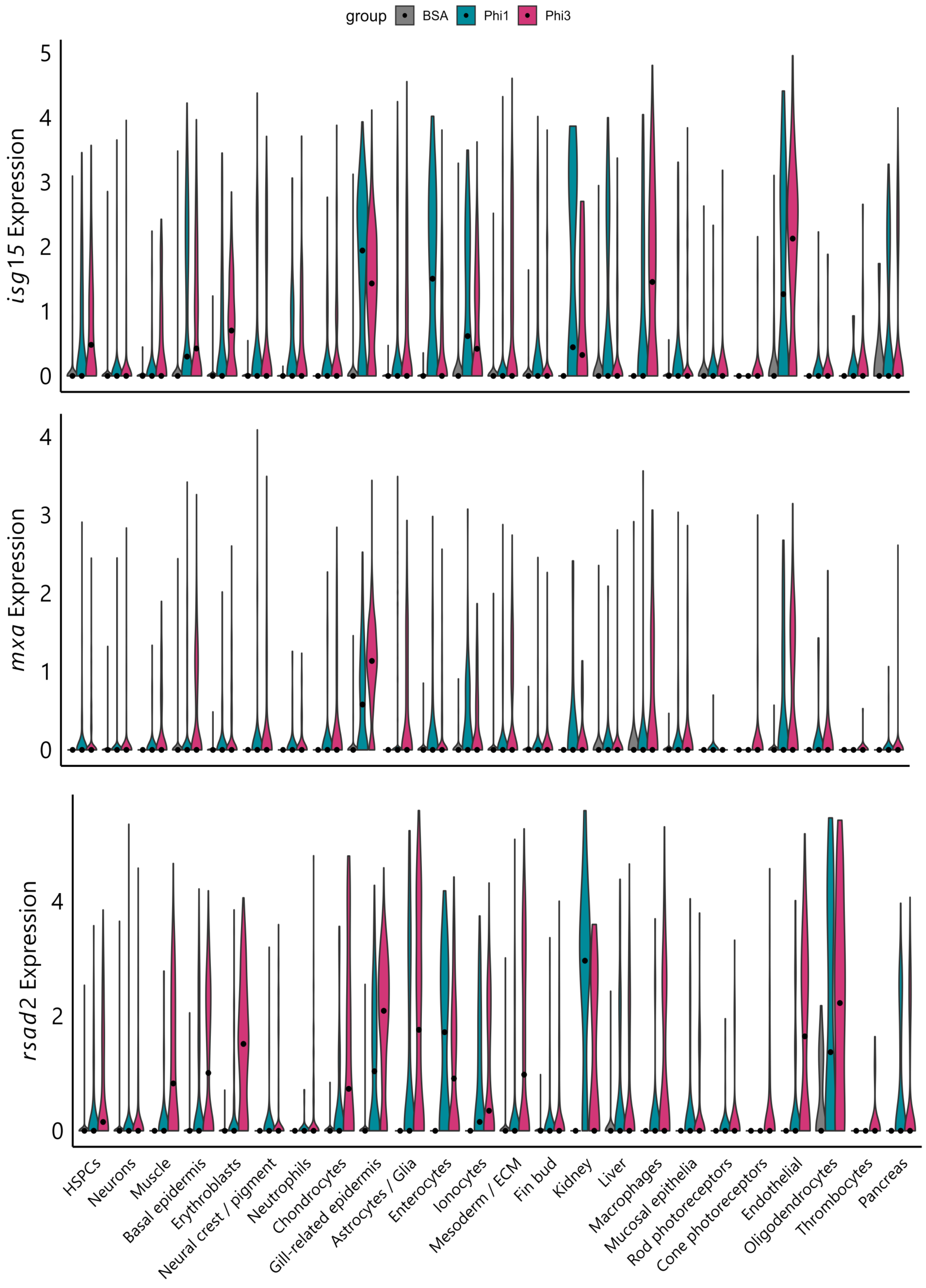


**Supplementary figure 4 – Violin plots of cell type expression of rsad2, isg15 and mxa**

Violin plots of log-normalised expression of *isg15*, *mxa*, and *rsad2* across annotated cell types for each treatment condition. Violins show distribution of expression within each cell type and the central dot represents the mean expression

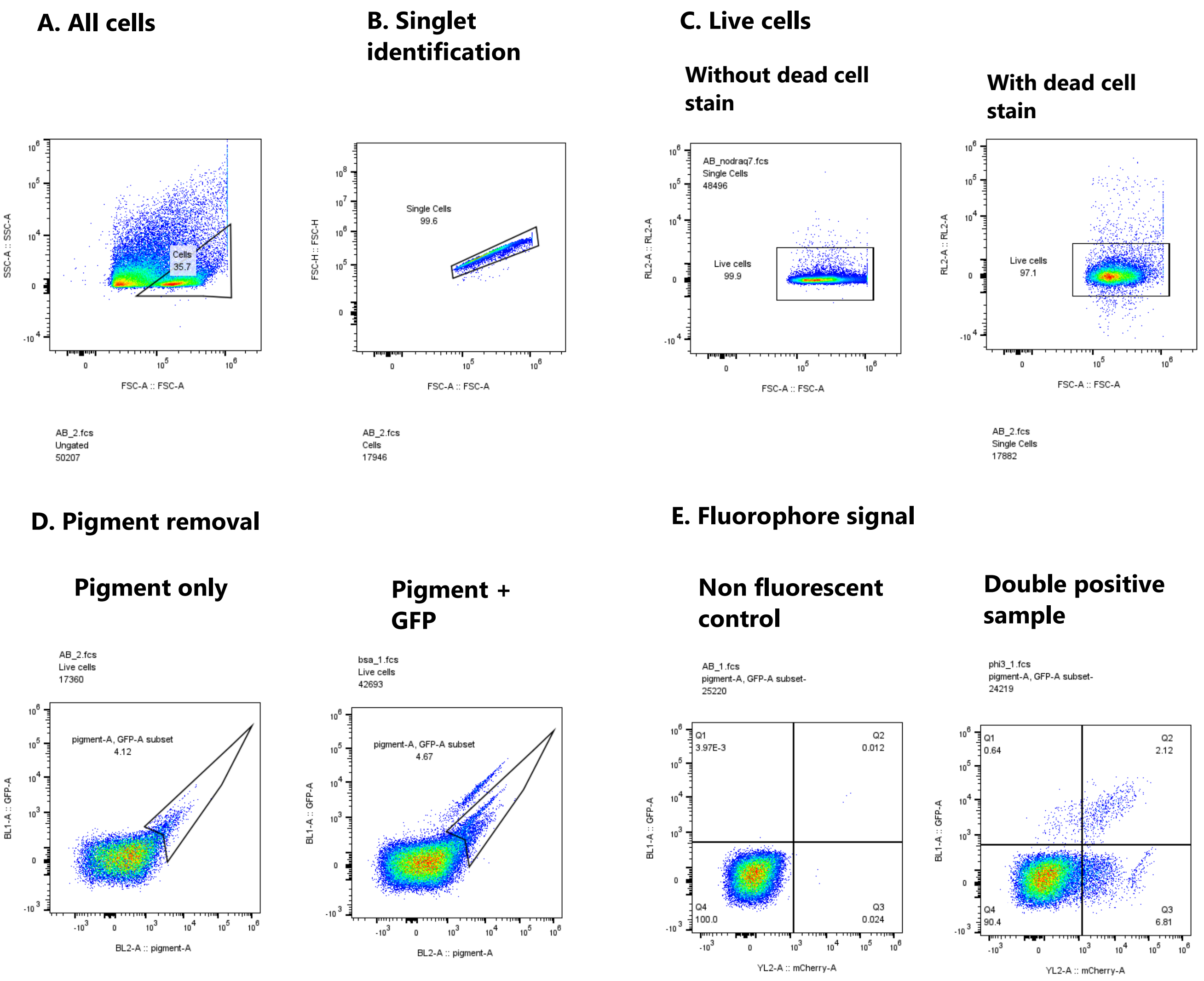


**Supplementary figure 5 – Gating strategy for the identification of mxa positive cells / enterocytes and macrophages**

**A.** Cells identified using FSC-A/SSC-A **B.** Singlets defined by FSC-H/FSC-A **C.** Live cells were defined based on gating of samples with and without the dead cell strain Draaq7 RL2-A/FSC-A. **D.** Autofluorescence from pigment cells removed by gating of pigmented non-fluorescent samples compared with GFP positive samples. **E.** Quadrants for GFP and mCherry positive cells were defined according
